# Establishment and validation of an endoplasmic reticulum stress reporter to monitor zebrafish ATF6 activity in development and disease

**DOI:** 10.1101/698811

**Authors:** Eric M. Clark, Hannah J.T. Nonarath, Jonathan R. Bostrom, Brian A. Link

**Affiliations:** Department of Cell Biology, Neurobiology and Anatomy, Medical College of Wisconsin, Milwaukee, WI, United States of America

**Keywords:** ATF6, endoplasmic reticulum stress, unfolded protein response, zebrafish, neurodegeneration

## Abstract

Induction of endoplasmic reticulum (ER) stress is associated with diverse developmental and degenerative diseases. Modified ER homeostasis causes activation of conserved stress pathways at the ER called the unfolded protein response (UPR). ATF6 is a transcription factor activated during ER stress as part of a coordinated UPR. ATF6 resides at the ER, and upon activation is transported to the Golgi apparatus where it is cleaved by proteases to create an amino-terminal cytoplasmic fragment (ATF6f). ATF6f translocates to the nucleus to activate transcriptional targets. Here, we describe establishment and validation of zebrafish reporter lines for *ATF6* activity. These transgenic lines are based on a defined and multimerized *ATF6* consensus site which drives either eGFP or destabilized eGFP (d2GFP), enabling dynamic study of *ATF6* activity during development and disease. The results show that the reporter is specific for the ATF6 pathway, active during development, and induced in disease models known to engage UPR. Specifically, during development, *ATF6* activity is highest in the lens, skeletal muscle, fins, and gills. The reporter is also activated by common chemical inducers of ER stress including tunicamycin, thapsigargin, and brefeldin A, as well as by heat shock. In both an ALS and a cone dystrophy model, *ATF6* reporter expression is induced in spinal cord interneurons or photoreceptors, respectively, suggesting a role for ATF6 response in multiple neurodegenerative diseases. Collectively our results show these *ATF6* reporters can be used to monitor *ATF6* activity changes throughout development and in zebrafish models of disease.

**Summary Statement:** We have established and validated transgenic zebrafish reporter lines to quantitatively measure the ATF6 branch of the endoplasmic reticulum stress pathway in development and disease.

## Introduction

The ER is an important organelle in the cell for biosynthesis, folding, and maturation of proteins destined for the cell membrane or the extracellular space (Görlach et al., 2006). The ER also plays other important roles including lipid biosynthesis (Sriburi et al., 2004), calcium homeostasis (Görlach et al., 2006) and genesis of autophagosomes (Hayashi-Nishino et al., 2009) and peroxisomes (Geuze et al., 2003). Resident to the ER are multiple chaperones, foldases, and cofactors to ensure rapid and functional protein production and to protect the ER when translational demand is elevated (Görlach et al., 2006). If the protein load becomes too high, the ER can expand to increase surface area (Schuck et al., 2009), or target misfolded proteins for degradation by the proteasome (ERAD) (Meusser et al., 2005; Stolz and Wolf, 2010) or the lysosome (Ishida and Nagata., 2009).

Another strategy to maintain ER homeostasis in the presence of misfolded proteins or other stress at the ER is through transcriptional regulation. This regulation can affect the described mechanisms as well as downregulate overall translation to decrease the protein quantity, while increasing the functional capacity of the ER to promote cell survival. If pro-survival modifications are insufficient, chronic ER stress can up-regulate transcription of pro-apoptosis machinery (Walter and Ron, 2011). These pro-survival and pro-apoptosis transcriptional modifications are downstream of three main UPR pathways, ATF6, IRE1, and PERK (Walter and Ron, 2011). In the normal state, ATF6, IRE1, and PERK proteins are tethered at the ER through interaction with Bip/Grp78 in the lumen. After ER stress, the tethered proteins are released, and able to activate UPR. The mechanisms for activation for each branch of the ER stress response network are conserved among vertebrates including mammals and fish (Ishikawa 2011).

ATF6 is synthesized as a 90KD protein with 670 amino acids. It is a member of the ATF/CREB basic-leucine zipper (bzip) DNA binding protein family. After ER stress-induced release of ATF6 from the ER, its Golgi localization signal is exposed. At the Golgi, ATF6 is cleaved by site 1 protease (S1P) and site 2 protease (S2P), leaving a 50KD cleavage product ATF6f (Hillary and FitzGerald, 2018). ATF6f translocates to the nucleus and can bind to ER stress-response elements (Roy and Lee 1999; Yoshida et al., 1998). Later, Wang et al., (2000) identified the specific consensus binding sequence for ATF6, TGACGTGGCGATTCC. In the current study, this consensus sequence was used to build a reporter for examining *ATF6* activation in vivo, which has not been previously monitored.

Recently, ATF6 has been implicated in development of multiple tissue types, and dysregulation of *ATF6* expression is related to disease (Hillary and FitzGerald, 2018). Developmental roles of ATF6 include mesoderm differentiation (Kroeger et al., 2018), osteogenesis and chondrogenesis (Guo et al., 2016; Jang et al., 2012; Kim et al., 2014; Maeda et al., 2015; Xiong et al., 2015), neurodevelopment (Cho et al., 2009; Naughton et al., 2015; Saito et al., 2012; Zhang et al., 2007) adipogenesis and lipogenesis (Lowe et al., 2012), and formation of female reproductive structures (Lin et al., 2012; Park et al., 2013; Xiong et al., 2016; Yang et al., 2015). Most relevant to our work is the role of ATF6 in ocular and muscular embryology, because these tissues had the highest *ATF6* reporter expression during zebrafish development. *ATF6* is dynamically expressed throughout lens formation (Firtina and Duncan, 2011). In muscle development, ATF6 acts in apoptosis (Nakanishi et al., 2005) and differentiation (Wang et al., 2015) processes. Dysregulation or loss of ATF6 during development and in adulthood is thought to contribute to neurological disorders. Recently, mutations in multiple domains of *ATF6* were identified in patients with autosomal recessive achromatopsia (Chiang et al., 2017; Kohl et al., 2015). The IRE1/XBP1 pathway has been implicated in retinitis pigmentosa (Chiang et al., 2015), suggesting that investigation of ATF6 and other UPR pathways is important in understanding photoreceptor homeostasis and disease. Multiple neurodegenerative diseases including amyotrophic lateral sclerosis (ALS) have misfolded proteins and signatures of ER stress. A *sod1 G93A* mouse model of ALS had elevated levels of Atf6f compared to wildtype mice at early symptomatic and end stages of disease suggesting that Atf6 could play an important role in progression of ALS and other neurodegenerative conditions (Kikuchi et al., 2006).

In zebrafish, Atf6 has been investigated in fatty liver disease and steatosis (Cinaroglu et al., 2011; Howarth et al., 2014), but how Atf6 is involved in other tissue types or disease processes has not been investigated. While cell stress reporters for monitoring *xbp1* splicing and Atf4 translational induction have been described in several species (Iwawaki et al., 2017; Kang et al., 2015; Li et al., 2015; Ryoo et al., 2013), there are currently no reporters for investigating *ATF6* activity in vivo for any animal model. To address this unmet need, we created a zebrafish reporter based on a previously identified *ATF6* consensus site (Wang et al., 2000) to drive either eGFP or destabilized eGFP, enabling dynamic study of *ATF6* activity during development and disease in vivo. *ATF6* was activated in multiple tissues throughout development, can be induced by ER stress, and is upregulated in zebrafish disease models of cone dystrophy and ALS. Together, our data show that the *ATF6* reporter is a useful tool for monitoring *ATF6* activity in real time and could be used in combination with other reporters to investigate the complicated nature of UPR signaling in development and disease progression.

## Results

### Establishment of an ATF6 response element reporter for in vivo monitoring of ER stress

Studies characterizing Atf6 activity in animals has relied on nuclear immunolocalization and expression analysis of target genes (Samali et al., 2010). These strategies have been employed in zebrafish, for example, to study liver biology (Cinaroglu et al., 2011; Howarth et al., 2014). However, there are no current strategies to directly measure *ATF6* activity in intact animals. Therefore, we generated transgenic reporters to quantitate *ATF6* transcriptional activity in zebrafish. The transgenes use a previously identified *ATF6* consensus binding site (Wang et al., 2000) multimerized five times upstream of a minimal *c-fos* promoter driving either *eGFP* or destabilized *eGFP* (*5XATF6RE*:GFP; Fig. 1A). By injecting the reporter plasmid into multiple zebrafish embryos, a consistent expression pattern was observed, demonstrating that regardless of the transgene insertion site, the activity pattern was consistent. *5XATF6RE*:d2GFP (Fig. 1B) and *5XATF6RE*:eGFP (Fig. 1C) transgene expression was observed highest in the lens and skeletal muscle. *5XATF6RE*:d2GFP expression was more dynamic, consistent with a higher turn-over rate through proteasomal targeting by a PEST domain. Skeletal muscle expression was highest at 1 day post fertilization (dpf) and decreased over time, whereas lens expression stayed relatively consistent with age. Furthermore, dynamic expression was also observed in the caudal and pectoral fins, gills, and brain through development (Fig. 1B). Interestingly, maternally inherited *5XATF6RE* derived eGFP protein resulted in ubiquitous fluorescence throughout embryonic development (Fig. 2). The destabilized transgenic version, *5XATF6RE*:d2GFP, did not show this maternal effect, suggesting the main source of fluorescence with the stable eGFP version is due to protein inheritance and perdurance, and not significant mRNA translation prior to zygotic expression. Because of the higher baseline fluorescence in embryos derived from maternal transgenes, paternally-provided transgenic embryos were analyzed in subsequent experiments.

**Figure 1.**
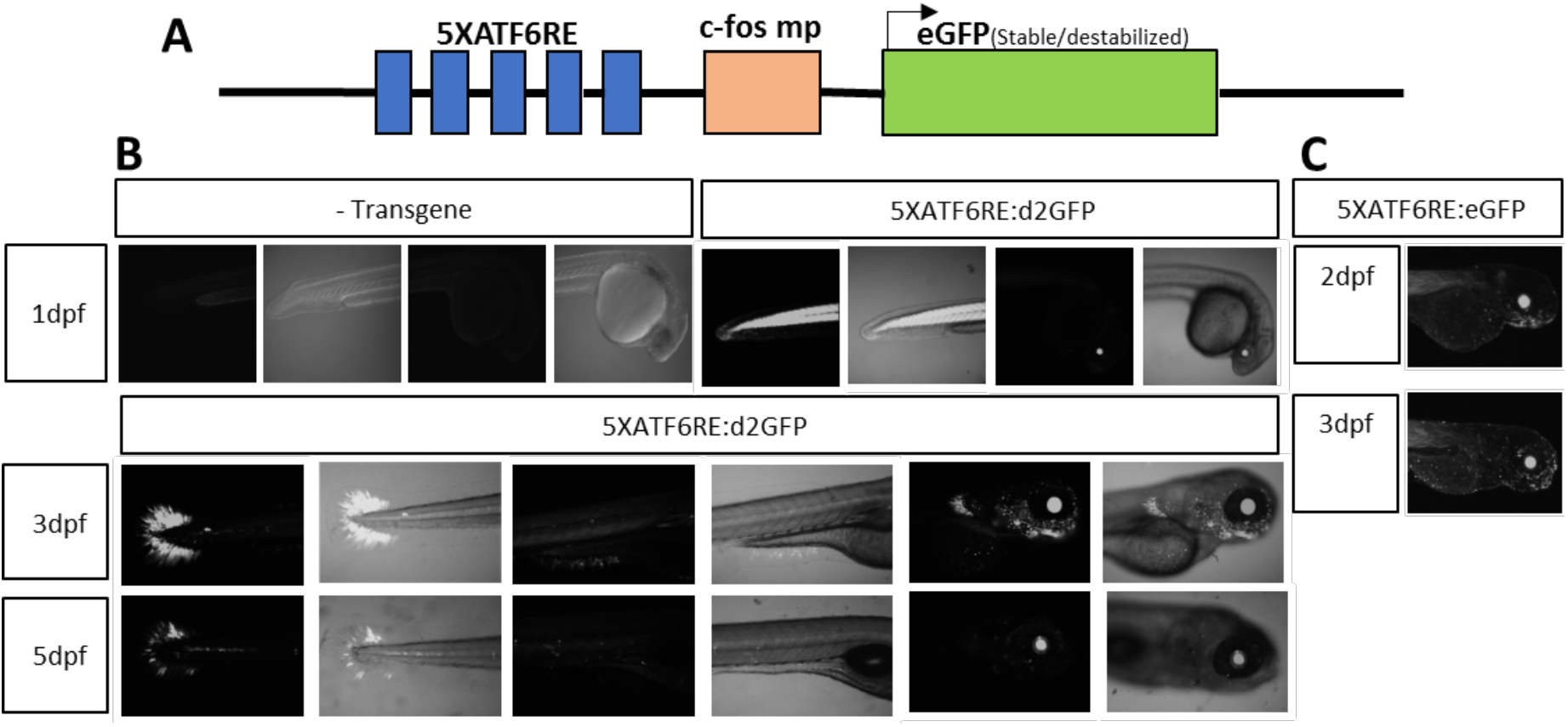
Reporter expression in developing zebrafish. (A) Schematic of the *ATF6* reporter plasmid showing 5 multimerized ATF6 binding sites (*5XATF6RE*) upstream of a *c-fos* minimal promoter (*c-fos mp*) driving expression of either destabilized GFP (*d2GFP*) or stable, enhanced GFP (*eGFP*). (B) Time-course showing *5XATF6RE*:d2GFP expression at 1,3, and 5dpf. (C) Time-course showing *5XATF6RE*:eGFP at 2 and 3 dpf.

**Figure 2.**
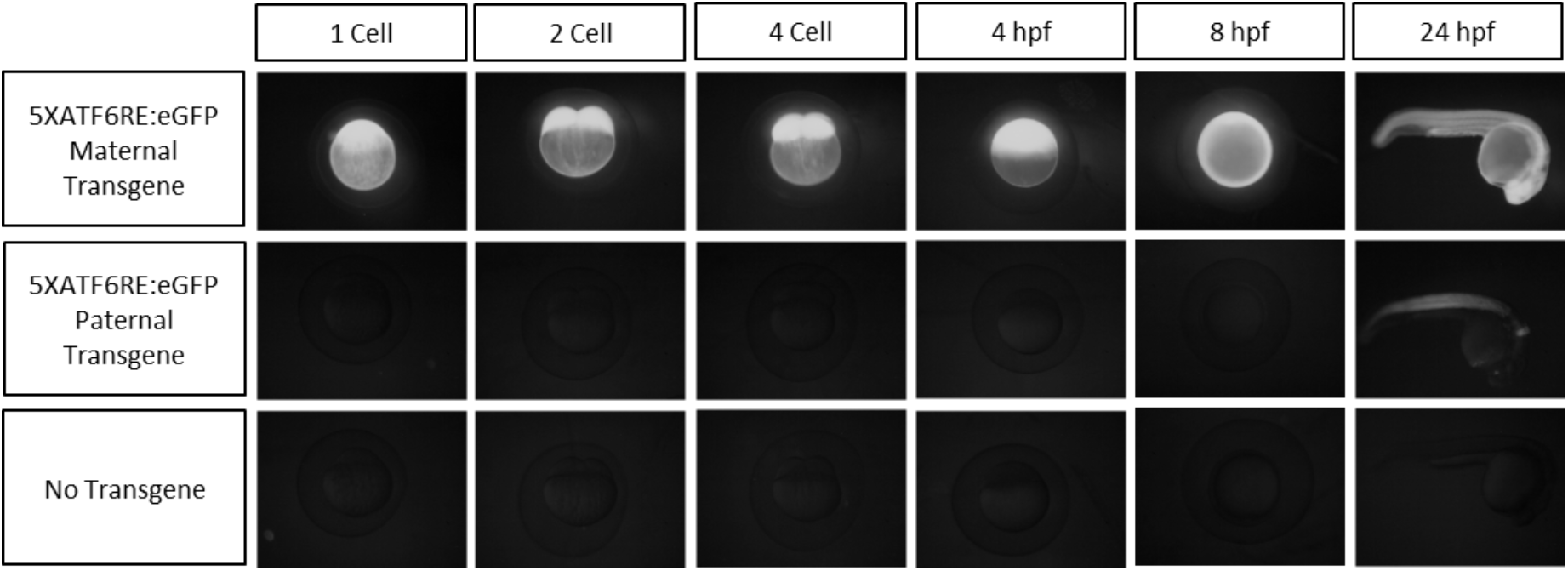
Maternal/paternal inheritance of eGFP from the *5XATF6RE*:eGFP transgene. Epifluorescent images showing embryos in which the *5XATF6RE*:eGFP was inherited maternally (Top) or paternally (Middle). Age-matched embryos not harboring a transgene are shown as an autoflourescence control (Bottom).

To quantitatively analyze *ATF6* reporter expression, the number of active transgene inserts was investigated. F1 zebrafish were outcrossed, and the number of F2 embryos with reporter expression were counted. With one copy of active transgene, approximately 50% of F2 embryos are expected to have reporter expression, which was observed in *5XATF6RE*:d2GFP and *5XATF6RE*:eGFP F2 embryos (Fig. S1A). Additionally, the lens expression at 2dpf was consistent between sibling embryos, demonstrating that endogenous expression of the transgene can be used for quantitative analysis (Fig. S1B, C).

### Characterization of transgene specificity

Previously, a consensus binding site was defined for ATF6 in HeLa cells (Wang et al., 2000). Here, we sought to confirm, and further investigate binding site specificity in zebrafish. Towards this goal zebrafish embryos expressing *5XATF6RE*:d2GFP and *hsp70*:GAL4 were injected with plasmids encoding active versions of transcription factors for each of the ER stress pathways (Fig. 3A). Zebrafish embryos were heat shocked at 2dpf to induce robust, but mosaic levels of each transcription factor. After imaging 12 hours post-heat shock, constitutive active ATF6 (*caATF6*) caused significantly higher *5XATF6RE*:d2GFP expression compared to the injection control (p=0.0067; Fig. 3B, C). Conversely, heat shock induced expression of spliced XBP1 (*XBP1s*) slightly, but significantly, inhibited *5XATF6RE*:d2GFP expression (p=0.0182; Fig. 3B, C), and *ATF4* had no significant effect (p=0.1055; Fig. 3B, C). Additionally, induced *caATF6* expression also activated the *5XATF6RE*:eGFP transgene (p=0.0071; Fig. S2). The *caATF6* expression resulted in autonomous transgene activation as cells expressing *caATF6-2A-mCherry* colocalized with *5xATF6RE*:d2GFP significantly more than mCherry-only expressing control cells (p=0.0005; Fig. S3). As 40% of d2GFP (ATF6 responsive cells) did not colocalize with mCherry (cells expressing caATF6), and 60% of mCherry-positive cells did not express detectible d2GFP, there is evidence for non-autonomous expression and non-responsive cells, respectively (Fig. S3). While ATF6 non-autonomy has not been previously described, Xbp1s has been shown to activate UPR in adjacent cells (Taylor et al., 2013; Williams et al., 2014).

**Figure 3.**
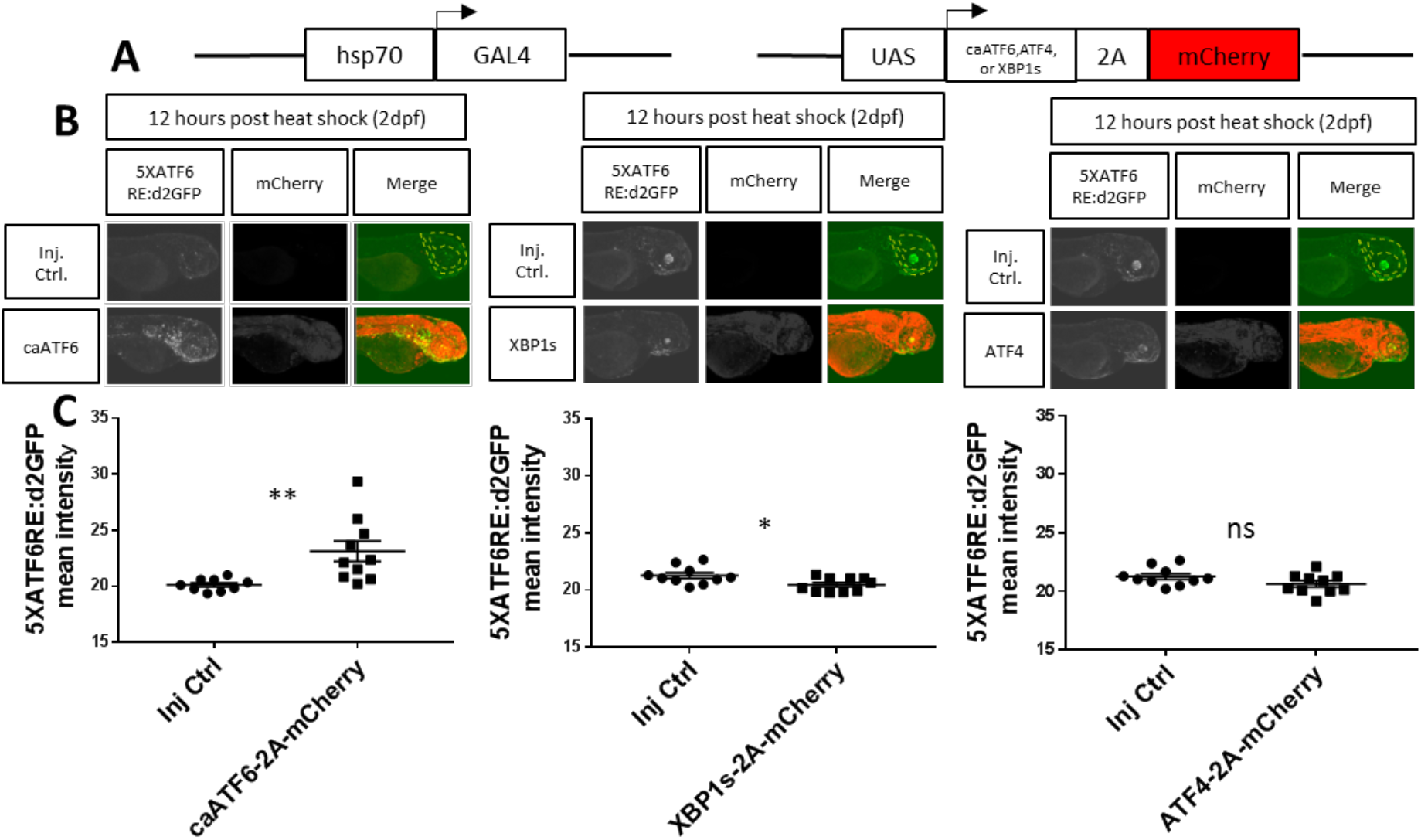
Heat shock induced constitutive active ATF6 specifically induces reporter expression. (A) Schematics of the *hsp70*:GAL4 construct used to induce ubiquitous expression of *UAS* constructs expressing either *caATF6, XBP1s, or ATF4* with a *T2A* self-cleaving peptide and *mCherry*. (BC) Representative images (B) and quantification of reporter expression (C) from embryos expressing *hsp70*:GAL4 and *5XATF6RE*:d2GFP transgenes that were injected with *mCherry* tagged *UAS* plasmids to overexpress *caATF6* (left), *XBP1s* (middle), or *ATF4* (right) compared to injection control (Inj. Ctrl.) embryos injected with all components except overexpression plasmids. 2dpf embryos were heat shocked and confocal images were captured 12 hours later. Quantification (dotted line) revealed a significant increase in *5XATF6RE*:d2GFP intensity with *caATF6* overexpression (p=0.0067), a significant decrease with *XBP1s* overexpression (p=0.0182), and no significant change with *ATF4* overexpression (p=0.1055) compared to the injection control. * p≤0.05, ** p≤0.01, ns p>0.05; unpaired t-test.

To further explore reporter specificity, we conducted loss of Atf6 function experiments. Two approaches were employed: Atf6 protein levels were diminished using an *atf6* translation blocking morpholino (Cingarolu et al., 2011), while Atf6 activity was inhibited by expression of a dominant negative ATF6 (*dnATF6*). Using these reagents, we tested whether *5XATF6RE*:d2GFP expression could be blocked following induction by tunicamycin. Tunicamycin inhibits N-linked glycosylation at the ER and is commonly used to induce UPR (Wang et al., 2000). We found injection of *atf6* morpholino significantly blocked tunicamycin induced *5XATF6RE*:eGFP expression (p=0.0150, Fig. 4A). Heat shock induced expression of *dnATF6* was variable, but the highest levels of dnATF6, as measured by mCherry intensity, completely negated *5XATF6RE*:d2GFP expression caused by tunicamycin treatment, resulting in a significant inverse correlation between dose of the dnATF6 and activation of the reporter (R^2^=0.4712, p=0.0284; Fig. 4B). Cumulatively, these gain and loss of function experiments indicate that *ATF6* reporter activity is specific for ATF6, and not responsive to other UPR pathways.

**Figure 4.**
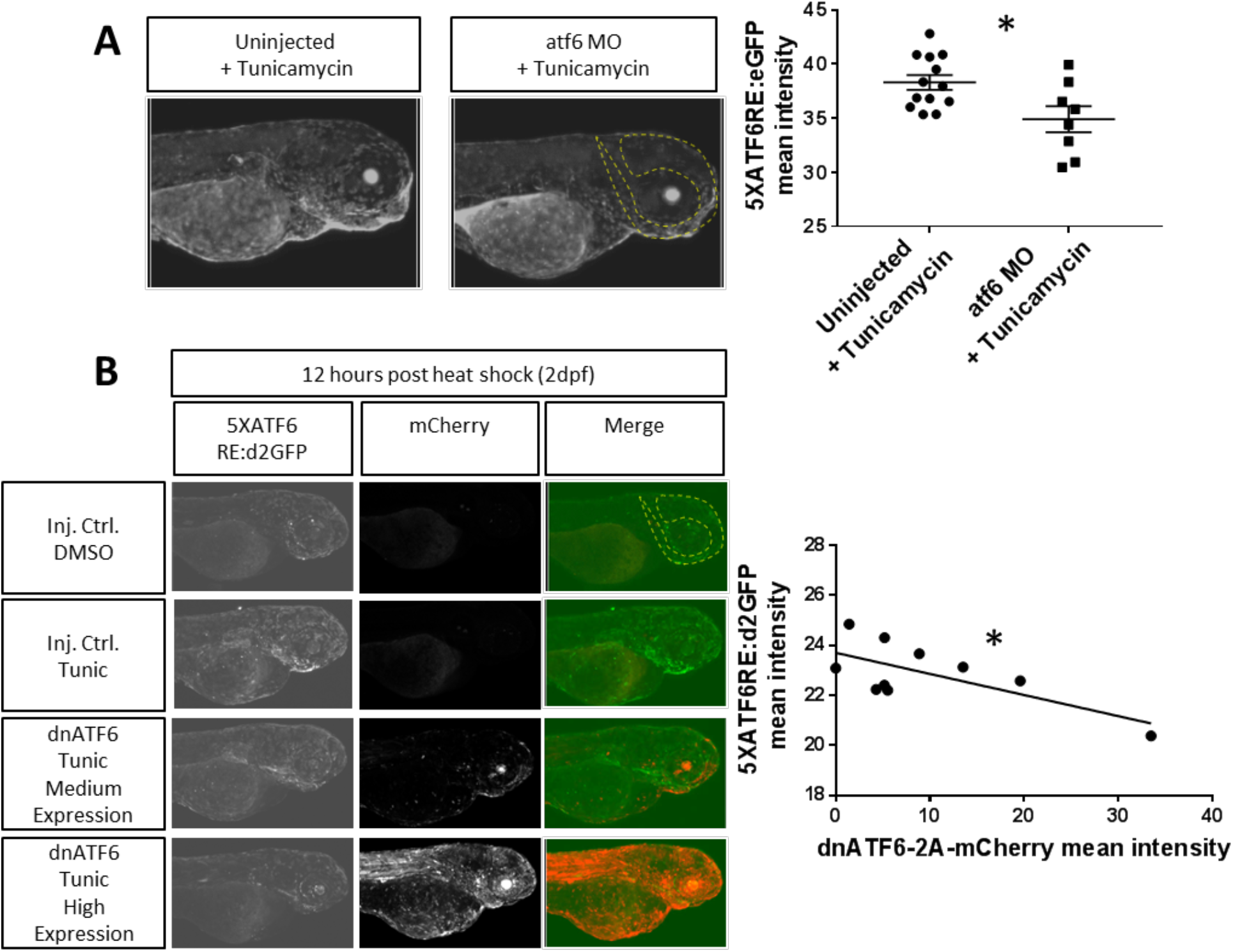
Tunicamycin induced reporter expression is blocked by inhibiting ATF6 translation and binding. (A) After injection of an *atf6* morpholino (*atf6* MO) at the 1-4 cell stage to inhibit ATF6 translation, 1dpf embryos were treated with Tunicamycin (Tunic) and expression at 3dpf in the head region (dotted line) was quantified. *atf6* MO injected embryos had significantly less *5XATF6RE*:eGFP expression compared to Tunicamycin only treated embryos (p=0.0150). Only embryos with detectable *5XATF6RE*:eGFP expression were analyzed. * p<.05; unpaired t-test (B) Embryos expressing *hsp70*:Gal4 and *5XATF6RE*:d2GFP transgenes were injected with an *mCherry* tagged *UAS* plasmid to overexpress dominant negative ATF6 (*dnATF6*) that inhibits endogenous ATF6 binding. 2dpf embryos were heat shocked, treated with Tunicamycin, and expression of GFP and mCherry was quantified (dotted line) 12 hours later. Representative images show that compared to a DMSO treated injection control, Tunicamycin treatment increases *5XATF6RE*:d2GFP expression, which is decreased with increasing dnATF6-2A-mCherry expression, resulting in a significant negative correlation (R^2^=0.4712, p=0.0284). *p≤0.05; Pearson’s correlation coefficient.

### Monitoring ATF6 reporter expression after induction of ER stress

The specific mechanism for how UPR is activated often results in differential pathway utilization. To probe how different chemical inducers of UPR affect *ATF6* activation, we bath applied compounds to transgenic zebrafish embryos. Zebrafish embryos respire through their skin until 7dpf (Rombough et al., 2002) and chemicals applied to the water are readily absorbed. After treatment of zebrafish embryos with a variety of established inducers of UPR, *5XATF6RE*:d2GFP expression was increased for most, but not all compounds as compared to DMSO treated control embryos (Fig. 5). Interestingly, for those activating the *ATF6* reporters, the dynamics and tissue responses were distinct for each chemical inducer of ER stress. Unexpectedly, Dithiothreitol (DTT) did not activate the *ATF6* reporter transgenes. DTT is a strong reducing agent that can denature proteins by preventing intra- and intermolecular disulfide bonds. In contrast to DTT mediated activation of *xbp1* (Li et al., 2015), DTT did not alter *5XATF6RE*:d2GFP expression as compared to DMSO treated control embryos (Fig. 5A). Tunicamycin, as noted previously, did induce reporter expression in a wide-spread manner, with peak levels at 48 hours post treatment (hpt). Thapsigargin is an inhibitor of the ER Ca2+ ATPase, resulting in calcium efflux and dysfunction of the organelle. Thapsigargin induced expression of *5XATF6RE*:d2GFP peaked 8 hours post treatment and subsequently decreased by 24 hpt (Fig. 5A). Brefeldin A inhibits vesicle formation at the Golgi and ultimately results in fusion of Golgi and ER membranes, leading to ER stress. Brefeldin A induced expression of *5XATF6RE*:d2GFP primarily in the yolk and gills, with a peak at 24 hpt (Fig. 5A). To further analyze differential *ATF6* activation in response to ER stressors, we more closely inspected spinal cord neurons (Fig. 5B). Treatment with Tunicamycin or Brefeldin A, but neither DMSO nor DTT, activated the *ATF6* reporter in a distinct subset of cells. Together, this analysis shows that the *ATF6* activity reporters respond to a variety of ER stressors but suggests that distinct mechanisms of ER insult results in differential response kinetics and tissue sensitivities. Like *5XATF6RE*:d2GFP, *5XATF6RE*:eGFP expression levels were also elevated after treatment with Tunicamycin, Thapsigargin, and Brefeldin A, but not DTT or DMSO (Fig. S4).

**Figure 5.**
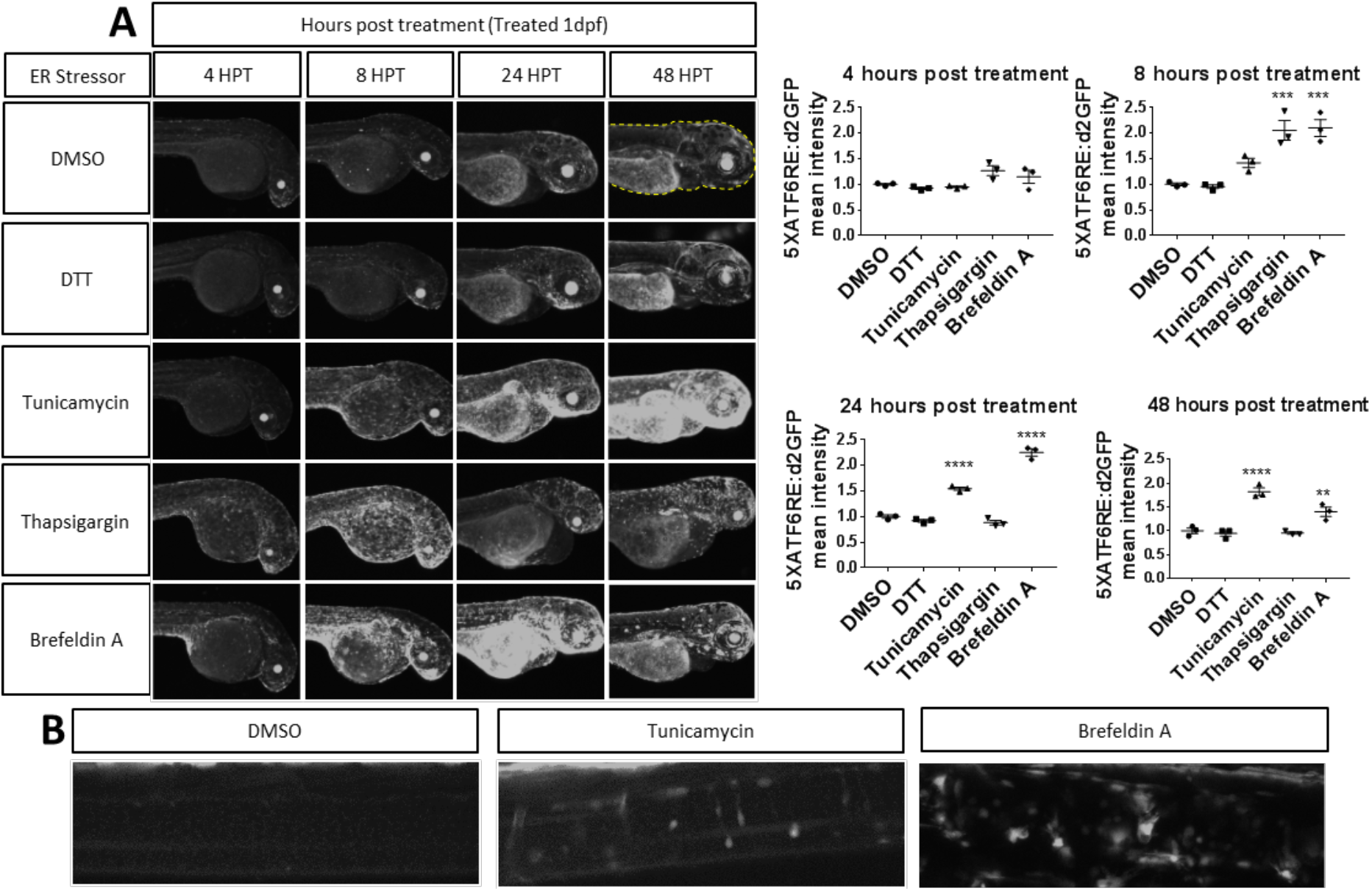
Chemical ER stressors activate reporter expression. (A) Zebrafish embryos were treated with ER stressors at 1dpf. Quantification (dotted line) revealed that *5XATF6RE*:d2GFP expression was significantly higher at 8hpt with Thapsigargin (p=0.0004) and Brefeldin A (p=0.0003) treatment, at 24hpt with Tunicamycin (p=0.0001) and Brefeldin A (p=0.0001) treatment, and at 48hpt with Tunicamycin (p=0.0001) and Brefeldin A (p=0.0065) treatment. ** p≤0.01; *** p≤0.001; **** p≤0.0001; unpaired one-way ANOVA with Dunnett’s post-hoc test with respect to DMSO control group. (B) Representative images showing increased *5XATF6RE*:d2GFP expression in the spinal cord of embryos 48hpt with Tunicamycin or Brefeldin A compared to the DMSO treated control.

In addition to chemical stress, heat stress can also be used to alter proteostasis (Long et al., 2012), and can induce apoptosis at 8 hours post-heat shock in the spinal cord (Yabu et al., 2001). We hypothesized that *ATF6* expression might also be induced by heat stress. Indeed, 8 hours after heat shock, *5XATF6RE*:d2GFP reporter expression was elevated in the spinal cord (Fig. 6A). Similarly, reporter expression was significantly higher in the lens (p<0.0001; Fig. 6B) and head region (p=0.0248; Fig. 6C) at various times post-heat shock. Therefore, *ATF6* transcriptional activity is elevated in multiple tissues in response to altered proteostasis.

**Figure 6.**
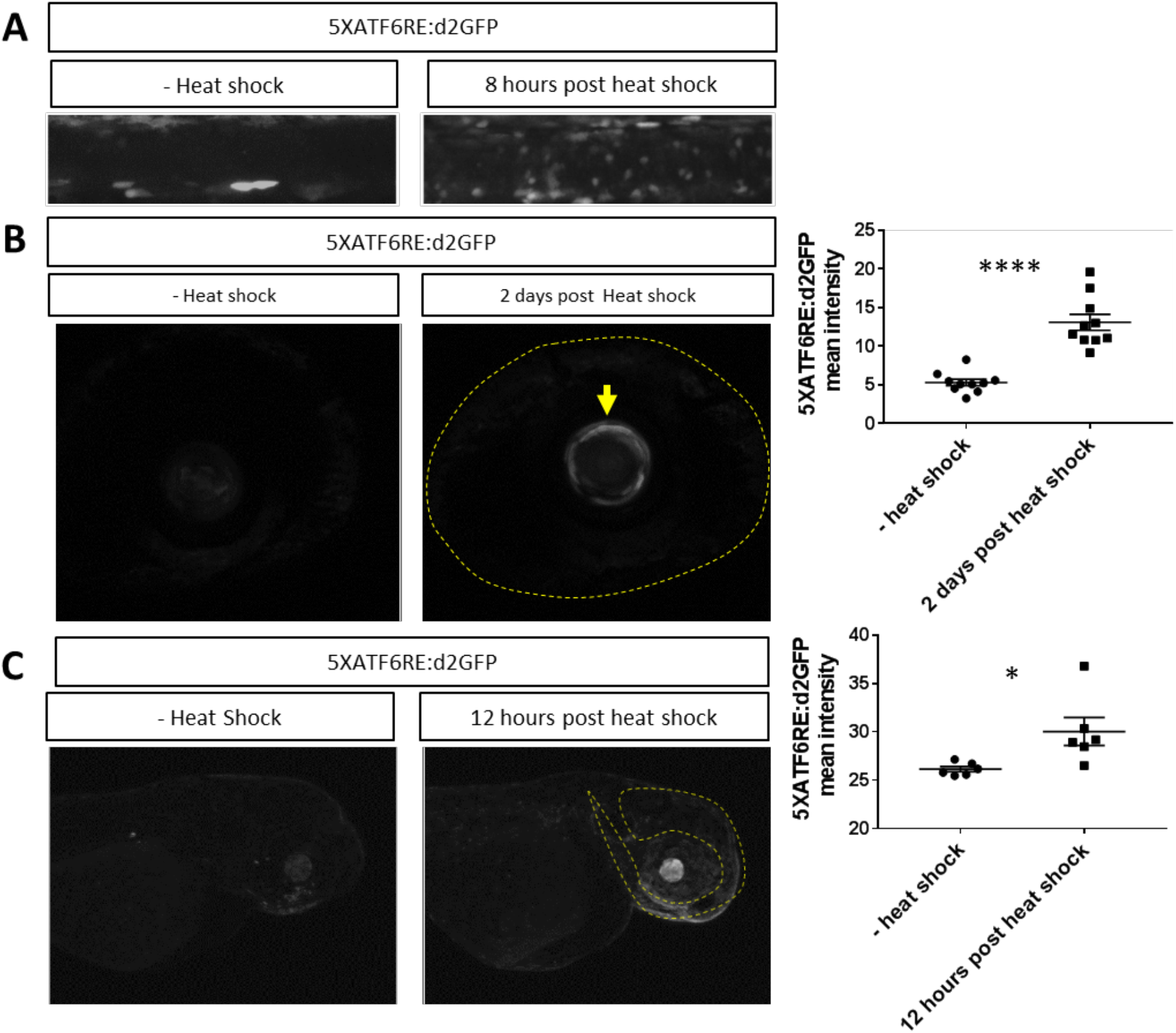
Heat shock activates reporter expression. (A) Representative images showing increased *5XATF6RE*:d2GFP expression in the spinal cord 8 hours after heat shock compared to controls without heat shock. (B) Representative images and quantification of the lens (arrow) revealed significantly higher levels of *5XATF6RE*:d2GFP expression 2 days post heat shock compared to controls without heat shock (p<0.0001). Dotted line outlines the eye. (C) Representative images and quantification of the head region (dotted lines) revealed significantly higher levels of *5XATF6RE*:d2GFP expression 12 hours post heat shock compared to controls without heat shock (p=0.0248). * p≤0.05; **** p≤0.0001; unpaired t-test.

### Investigation of ATF6 in neurodegenerative disease

The results above suggest that eye and spinal cord tissue may be susceptible to altered proteostasis and ER stress. Many neurodegenerative diseases are characterized by misfolded proteins and UPR activation. Neurons, including photoreceptors, are highly metabolically active with a significant protein turn-over demand, and therefore could be more susceptible to ER stress. In fact, photoreceptors are the most metabolically active cell type in the human body (Sung and Chuang, 2010; Wong-Riley, 2010). Interestingly, in mice and humans, mutation to *ATF6* results in age-dependent photoreceptor degeneration (Ansar et al., 2015; Kohl et al., 2015). This result suggests an important protective effect of ATF6 in these photoreceptor cells. Therefore, we hypothesized that ATF6 signaling is responsive to misfolded proteins that accumulate at the ER in photoreceptors. When a mis-folding prone mutant cone opsin (OPN1MW^W177R^), which is retained in the ER (Gardner et al., 2010), was expressed in cone photoreceptors, there was a dramatic increase in *ATF6* reporter expression (Fig. 7A). These findings support previous work that ATF6 signaling is important for photoreceptor homeostasis and demonstrate the utility of the *ATF6* activity reporters in studying neurodegenerative disease.

**Figure 7.**
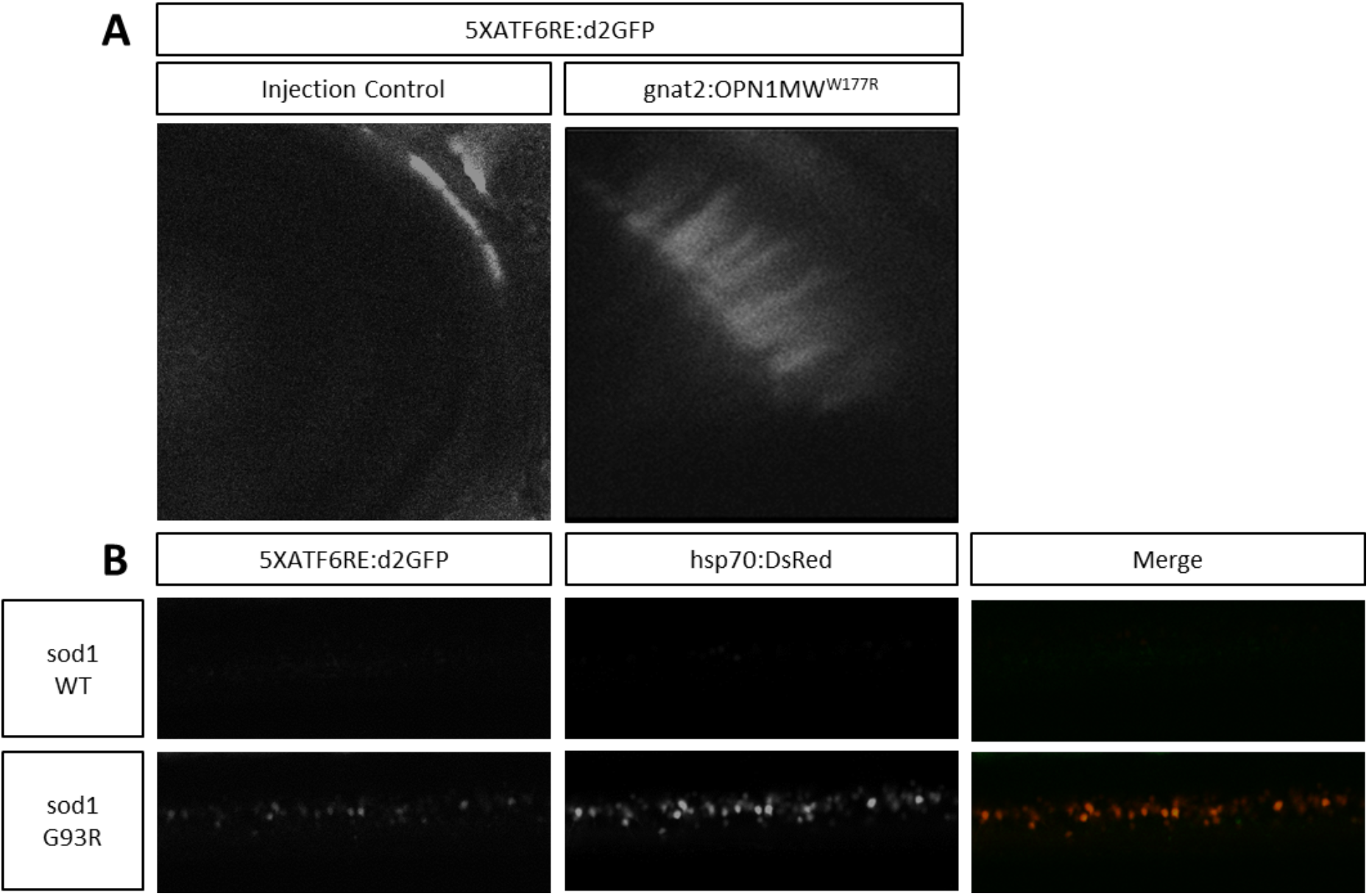
Reporter expression is activated in ALS and cone dystrophy zebrafish disease models. (A) Representative images showing that embryos injected with *gnat2*:OPN1MW^W177R^ mutant opsin have elevated *5XATF6RE*:d2GFP expression in photoreceptors which is absent in injection control embryos injected with all components except the overexpression construct. (B) Representative images showing overlap of *hsp70*:DsRed and *5XATF6RE*:d2GFP expression in *sod1 G93R* mutant spinal cord, which is absent in *sod1 WT* control spinal cord.

Amyotrophic lateral sclerosis (ALS) is another neurodegenerative disease characterized by misfolded proteins. Although a diverse group of genes result in ALS when mutated, a common pathology includes protein aggregation, activation of UPR, and death of motor neurons (Matus et al., 2013). One of the most common genes mutated is superoxide dismutase (*SOD1*). Defects in SOD1 that result in ALS are not due to loss of redox control, but instead are due to toxic gain of function that increases the rate of protein misfolding (Bruijn et al., 1998; Wiedau-Pazos et al. 1996; Yim et al., 1996). Motor neurons with mutations to amino acid G93 of *SOD1* are particularly susceptible to misfolding (Stathopulos et al., 2003). Based on this observation, a zebrafish model of ALS was established by transgenic overexpression of *sod1 G93R* (McGown et al., 2013; Ramesh et al., 2010). Using a general cell stress reporter gene (*hsp70*:DsRed), McGown and colleagues demonstrated that mutant *sod1 G93R* zebrafish embryos had elevated stress in the spinal cord, which did not occur in transgenic zebrafish expressing *WT sod1* (McGown et al., 2013). Interestingly, these cells were identified as interneurons, mostly glycinergic interneurons, suggesting that this cell type might be particularly susceptible in disease progression at pre-symptomatic stages of ALS. ATF6 is elevated in *sod1 G93A* mice and in human ALS patients (Atkin et al., 2006; Atkin et al., 2008; Prell et al., 2019). Therefore, we tested the role of ATF6 in an ALS zebrafish model expressing the *hsp70*:DsRed reporter. *5XATF6RE*:d2GFP colocalized with *hsp70*:DsRed expression in *sod1 G93R* zebrafish spinal cords, but both stress reporters were absent in *sod1 WT* zebrafish (Fig. 7B). These results indicate that multiple protective mechanisms including chaperone expression and UPR are increased in stressed interneurons early in ALS disease progression. Ultimately, these observations suggest that neurons are susceptible to unfolded protein stress at the ER, and that ATF6 could be an important target in modifying neurological diseases.

## Discussion

Investigation of Atf6 dynamics in whole animals has been limited to monitoring Atf6 immunoreactivity or target gene expression. Furthermore, in zebrafish, detailed studies of Atf6 have been conducted on the liver, but its role in other tissues has not been well characterized (Cinaroglu et al., 2011; Howarth et al., 2014). The development of transgenic zebrafish that report *ATF6* activity will facilitate studies across tissues and over time. Here we describe establishment and validation of *ATF6*-responsive transgenic zebrafish. The transgenes demonstrate dynamic activation during development and the transgenic fish can be used to monitor *ATF6* activity in diseases characterized by ER stress and unfolded proteins. In development, *ATF6* reporters were expressed highest in the lens and skeletal muscle but were also present in fins, central nervous system, and the branchial arch region. Elevated *ATF6* activity in the lens and muscle is consistent with those tissues expressing high levels of Atf6 during development (Firtina and Duncan, 2011; Nakanishi et al. 2005; Wang et al., 2015). Similar to maternal XBP1Δ-GFP expression (Li et al., 2015), maternal *5XATF6RE*:eGFP expression was pronounced, suggesting increased demand on ER function during oogenesis and/or embryogenesis immediately following fertilization and thus a requirement for stress protection.

With regard to disease relevance, we showed *ATF6* transgenes are responsive to cone photoreceptor expression of a variant opsin protein (OPN1MW^W177R^) that is known to misfold and cause photoreceptor degeneration in humans (Gardner et al., 2010). While ER retention has been characterized with the mutant cone opsin, the cell stress-response had not been previously characterized. Photoreceptors may be particularly sensitive to ER stress, as they synthesize and traffic tremendous amounts of phototransductive machinery as part of the daily shedding of their outer segments (Pearring et al., 2013). Supporting this idea are several observations. First, intravitreal injection of tunicamycin, a potent ER stress inducer and activator of *ATF6* as defined in our initial studies, results in severe photoreceptor degeneration (Alavi et al., 2015; Shimazawa et al., 2007). Furthermore, light induced retina degeneration also shows signs of ER stress, suggesting UPR may be a common pathway activated in photoreceptor degeneration (Kroeger et al., 2012). Perhaps most significantly, mutations to *ATF6* cause achromatopsia and result in loss of cone photoreceptor cells (Kohl et al., 2015). Characterization of a wide array of *ATF6* mutant alleles suggests patients have elevated risk to ER-stress that is compounded by both ER-retention of the mutant protein and the absence of ATF6 function (Chiang et al., 2017). ATF6 is also known to be activated during ALS and other neurodegenerative diseases. For example, *ATF6* is activated in the spinal cord of ALS patients (Atkin et al., 2008), and *ATF6* expression is decreased following disruption of the ALS causing *VAPB* gene (Chen et al., 2010; Nardo et al., 2011). Using a zebrafish model of ALS in which a disease allele of *sod1* (G93R) is expressed from the *sod1* promoter (Ramesh et al., 2010), we found elevated *ATF6* activity in spinal cord interneurons. These observations in both photoreceptors and spinal neurons, suggest that modifying *ATF6* activity could be a therapeutic approach to mitigate neurodegenerative diseases. Indeed, de-repression of Atf6 in a mouse model of Huntington’s disease provides neuroprotection to both striatal and hippocampal neurons (López-Hurtado et al., 2018; Naranjo et al., 2016). Furthermore, a small molecule activator of ATF6 reduces amyloidogenic protein secretion and aggregation, providing additional justification for targeting the ATF6 pathway in neurodegenerative disease (Plate et al., 2016).

One unexpected observation from our experiments was that, in addition to autonomous activation of the ATF6 activity reporter (*5XATF6RE*:d2GFP) by expressing ATF6 protein (*caATF6-2A-mCherry*), there was also non-autonomous reporter response. Cells proximal to those expressing constitutively activated ATF6 were often marked by high d2GFP expression without detectible *caATF6-2A-mCherry* expression. We performed a time-course of d2GFP expression at 3, 6, 9, and 12 hours following induction of *caATF6-2A-mCherry*, but never measured complete overlap in expression of mCherry and d2GFP, suggesting that the non-autonomy is not due to differential fluorescence turnover of either marker gene (data not shown). These findings are interesting in the context of work in *C. elegans* demonstrating that ER stress within neurons triggers autonomous *xbp1* activation that results in release of small clear vesicles. Signals packaged within the vesicles appear to trigger Xbp1 mediated UPR non-autonomously within peripheral tissues, which ultimately promotes stress-resistant longevity for the entire worm (Taylor et al., 2013). Similarly, expression of *xbp1s* in mouse hippocampal neurons, leads to non-autonomous activation of ER stress in the liver (Williams et al., 2014). More detailed investigation into the mechanisms underlying ATF6 non-autonomy in zebrafish could be accomplished with tools described here.

As an experimental resource to study homeostasis and disease, there are multiple potential uses for the *ATF6* reporter lines. Although small molecule screens have recently identified activators (Plate et al., 2016) and inhibitors (Gallagher and Walter, 2016) of the ATF6 pathway, additional targets of the pathway may prove beneficial. Zebrafish are particularly amenable for chemical-genetic screens and having whole animal readouts for *ATF6* activity provides a sophisticated platform for such assays. In addition to screens, the reporter lines will facilitate analysis of cells actively responding to ER stress. For example, by sorting cells based on their fluorescence, transcriptomics, metabolomics or proteomics could reveal susceptible cell types and cell states. In addition, such analysis might also reveal other pathways co-activated / inhibited with *ATF6* induction. These and certainly other studies will shed more light on the mechanism and potential modulation of ATF6 signaling.

## Materials and Methods

### Fish maintenance

Zebrafish (*Danio rerio*) were maintained at 28.5°C on an Aquatic Habitats recirculating filtered water system (Aquatic Habitats, Apopka, FL) in reverse-osmosis purified water supplemented with Instant Ocean salts (60mg/l) on a 14 h light: 10 h dark lighting cycle and fed a standard diet (Westerfield, 1995). All animal husbandry and experiments were approved and conducted in accordance with the guidelines set forth by the Institutional Animal Care and Use Committee of the Medical College of Wisconsin.

### Generation of plasmids

*ATF6RE*:eGFP and *ATF6RE*:d2GFP reporter plasmids with a *c-fos* minimal promoter were amplified by Genescript from Adgene plasmid 11976 (Wang et al., 2000), which contains the *ATF6* response element TGACGTGGCGATTCC interposed by linker sequences and multimerized 5 times (*5XATF6RE*). The purified *5XATF6RE* was placed into a 5’ entry clone and the three-part Gateway system (Thermo Fisher Scientific, Waltham, MA) was used to assemble the 5’ *5xATF6* response element in front of a *c-fos* minimal promoter followed by either *eGFP* or *d2GFP* and a 3’ *polyA tail*. The backbone vector containing Tol2-inverted repeats flanking the transgene constructs was used to facilitate plasmid insertion into the zebrafish genome (Fig. 1A; Kawakami et al., 2000; Kwan et al., 2007). Similarly, middle entry plasmids used for overexpression experiments were made by PCR amplification from parent plasmids, followed by Gateway recombineering using an upstream-activator sequence (*UAS*) promoter and a downstream *T2A* self cleaving peptide fused to *mCherry*, to mark expressing cells. Parent and final plasmids are listed in Table S1. Overexpression experiments using each plasmid was accomplished via the *GAL4/UAS* system (Scheer and Campos-Ortega, 1999) using *hsp70*:GAL4 expressing zebrafish for inducible, ubiquitous expression after heat shock at 39°C for 1 hour. For experiments, embryos were injected at the 1-4 cell stage with 9.2 nl of working solution containing 10ng/ul construct and 5ng/ul transposase. To construct the OPN1MW^W177R^ plasmid, the OPN1MW^W177R^ sequence was PCR amplified, inserted into a middle entry vector, and then added to a Tol2 destination vector downstream of a cone photoreceptor cell specific promoter (*gnat2*) using the gateway system.

### Microinjection and generation of transgenic ATF6 reporter zebrafish

Transposase mRNA was injected with *ATF6RE*:eGFP or *ATF6RE*:d2GFP plasmid DNA to generate F0 transgenic lines. F0 fish were analyzed to ensure consistent transgene expression regardless of the integration site. F0 Injected fish were raised to adulthood and outcrossed to wildtype zebrafish to establish F1 ATF6 reporter zebrafish with germline integration of the transgene. F1 transgenic larvae were raised to adulthood and outcrossed to wildtype zebrafish for expression analysis (Fig. S1). F2 zebrafish transgenic lines containing 1 active copy of each transgene were used for all subsequent analysis. These lines are designated Tg(Hsa.*ATF6RE*:eGFP)mw84 or Tg(Hsa.*ATF6RE*:d2GFP)mw85.

### Chemical and heat shock stress in zebrafish embryos

Chemical induction of ER stress in zebrafish embryos was accomplished by using a final concentration of .5mM DTT (NEB #B1034S; Invitrogen Y00147), 1uM Thapsigargin (Sigma #T9033), 1ug/ml Tunicamycin (Sigma #T7765), and 5uM Brefeldin A (Sigma #B7651) in 10ml PTU in a 16mm X 50mm petri dish. For heat stress, petri dishes containing embryos in 5mLs PTU and sealed using parafilm were incubated in a water bath set to 39°C for 1 hour.

### Morpholinos

*atf6* ATG morpholino, 5’-ACATTAAATTCGACGACATTGTGCC-3’ (Cinaroglu et al., 2011), was synthesized by GeneTools, LLC. 9.2nL of a working solution containing 700uM morpholino was injected into zebrafish embryos at the 1-4 cell stage.

### Image acquisition and analysis

For maternal contribution experiments, a Leica MZ FLIII epifluorescent stereomicroscope was used for image acquisition. *ATF6* reporter activity was observed with a Nikon Eclipse E600FN confocal system (Nikon, Tokyo, Japan) using the Nikon EZ-C1 software and a 10X air, .3NA or a 40X water, .8NA objective (Nikon Instruments). Reporter expression was quantified using ImageJ (Rasband, W.S., ImageJ, U.S. National Institutes of Health, MD). For tissue specific quantification, a region of interest (ROI) was drawn around the site of interest. A maximum pixel intensity projection image from a z-stack containing the total fluorescence in the ROI was analyzed for mean pixel intensity. For colocalization analysis Imaris 3D coloc software was used (BitPlane).

### Statistical analysis

Mean pixel intensity measurements of reporter expression were processed using Microsoft Excel (Microsoft, Redmond, WA) and graphed using GraphPad Prism (GraphPad, La Jolla, CA). An unpaired, two-tailed t test was used to analyze graphs with two groups. For three or more groups, a one-way ANOVA was conducted with Dunnett’s post-hoc analysis for pair-wise comparisons.

## Acknowledgements

The authors thank Michael Cliff, William Hudzinski, and Edi Kuhn for zebrafish husbandry. We also thank Dr. Christine Beattie for sharing *sod1* transgenic lines. We are also grateful to Dr. Alison Hardcastle for sharing the OPN1MW^W177R^ plasmid.

## Author contributions statement

E.M.C., H.J.T.N and B.A.L. designed experiments. E.M.C, H.J.T.N, and J.R.B performed experiments. E.M.C. analyzed experiments. E.M.C. and B.A.L wrote the manuscript. All authors read and approved the final manuscript.

---

**Figure S1.**
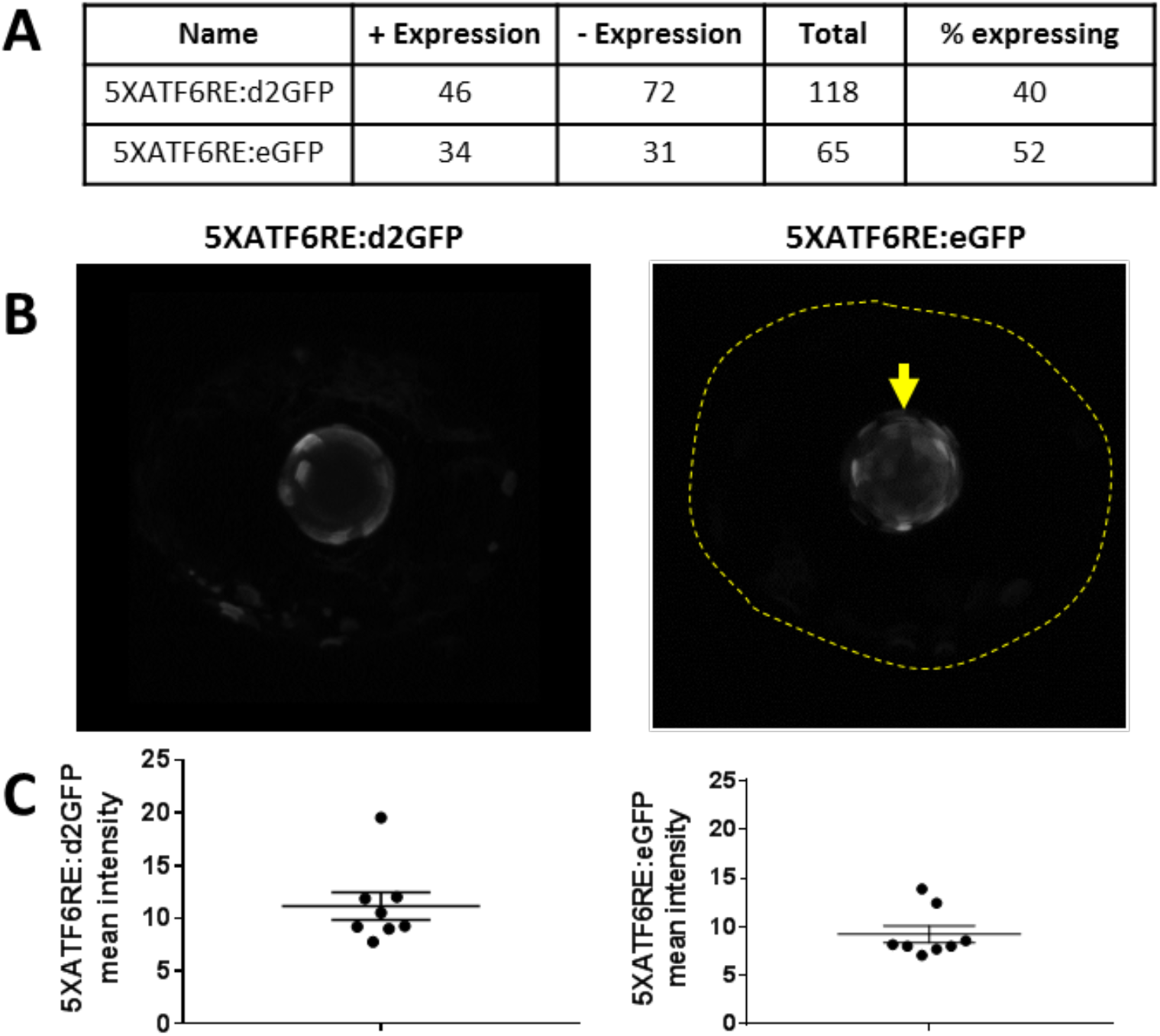
Single active transgene expression is consistent across embryos. (A) Number of F2 embryos expressing *5XATF6RE*:d2GFP or *5XATF6RE*:eGFP after an F1 outcross. (B,C) Representative images (B) and quantification (C) of reporter expression in the lens (arrow) from 2dpf F2 generation embryos from an F1 outcross. The dotted line outlines the eye.

**Figure S2.**
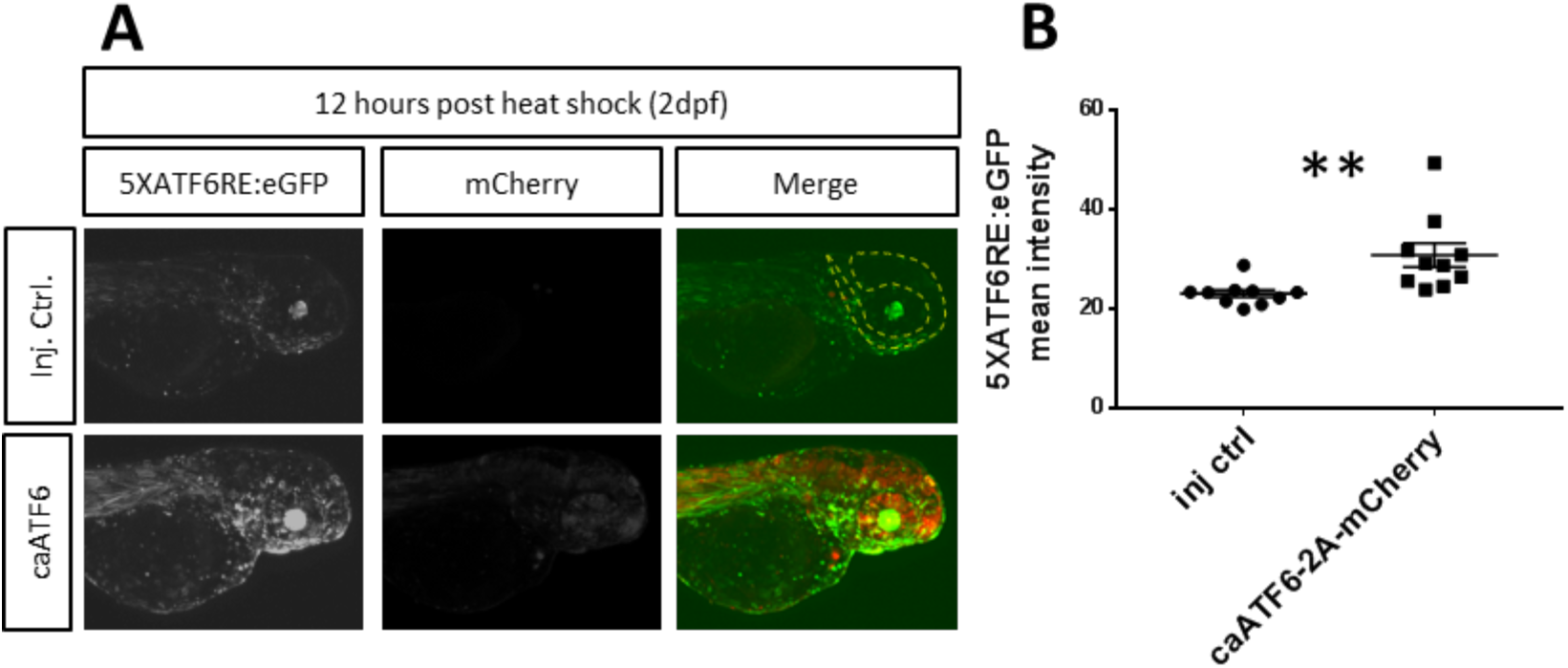
eGFP reporter expression is increased after heat shock induced constitutive active ATF6 expression. (AB) Representative images (A) and quantification of reporter expression (B) from embryos expressing *hsp70*:GAL4 and *5XATF6RE*:eGFP transgenes that were injected with an *mCherry* tagged *UAS* construct to overexpress *caATF6* compared to injection control (Inj. Ctrl.) embryos injected with all components except overexpression constructs. 2dpf embryos were heat shocked and confocal images were captured 12 hours later. Quantification (dotted line) revealed a significant increase in *5XATF6RE*:eGFP expression with overexpression of *caATF6* compared to the injection control (p=0.0071). ** p≤0.01; unpaired t-test.

**Figure S3.**
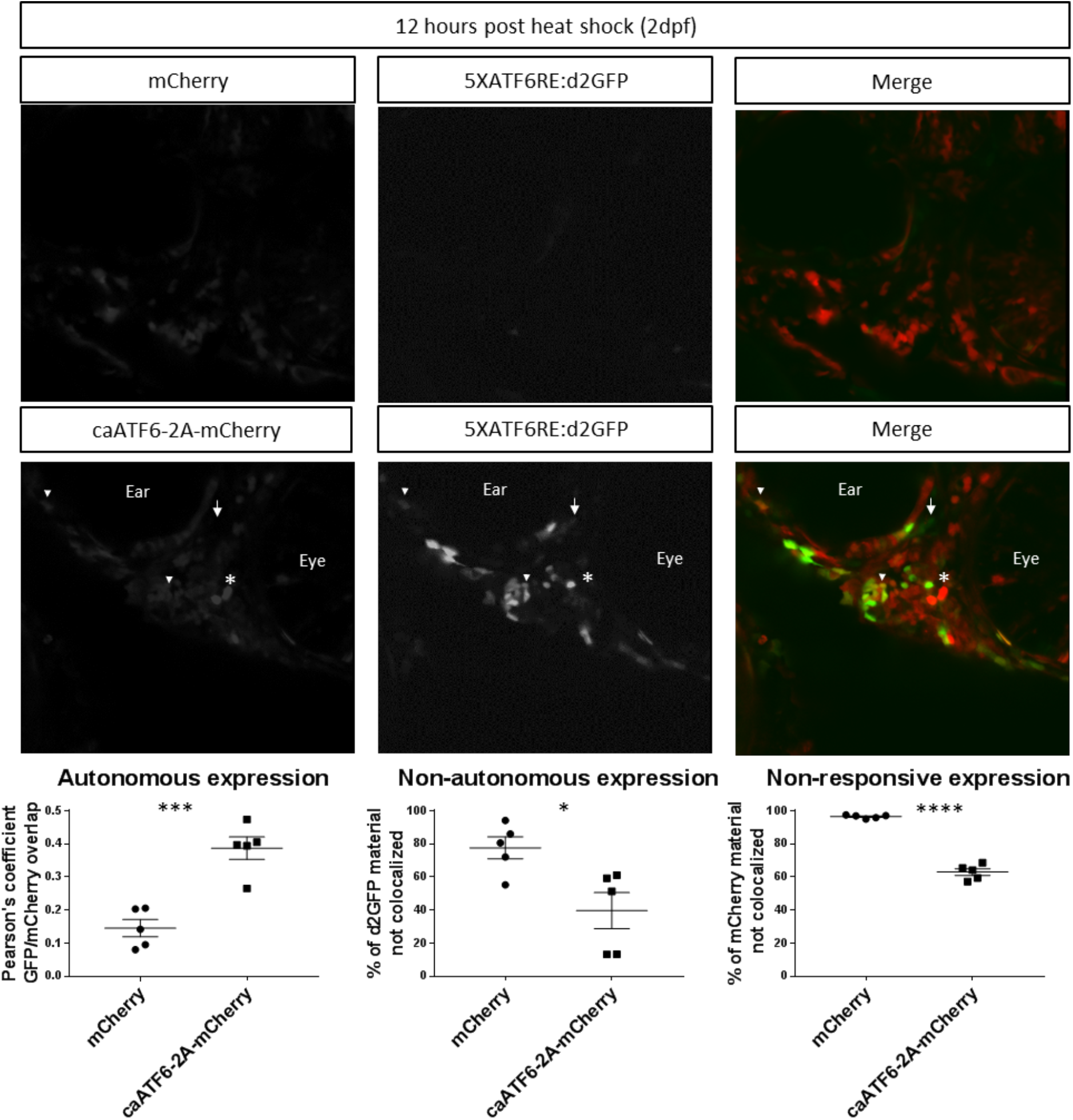
Autonomous reporter activation by constitutive active ATF6. Embryos expressing *hsp70*:GAL4 and *5XATF6RE*:d2GFP transgenes were injected with an *mCherry* tagged *UAS* construct to overexpress *caATF6* or a *UAS* construct with *mCherry* expression alone as a control. 2dpf embryos were heat shocked and confocal images zoomed on the jaw region where reporter expression was highest were captured 12 hours later. Quantification revealed that compared to *mCherry* expression alone, caATF6-2A-mCherry has significantly higher autonomous expression (p=.0005; arrowheads) and significantly lower non-autonomous expression (p=0.0177; arrow) and non-responsive expression. (p<0.0001; asterisk). * p≤0.05; ***p≤0.001; ****p≤0.0001; unpaired t-test.

**Figure S4.**
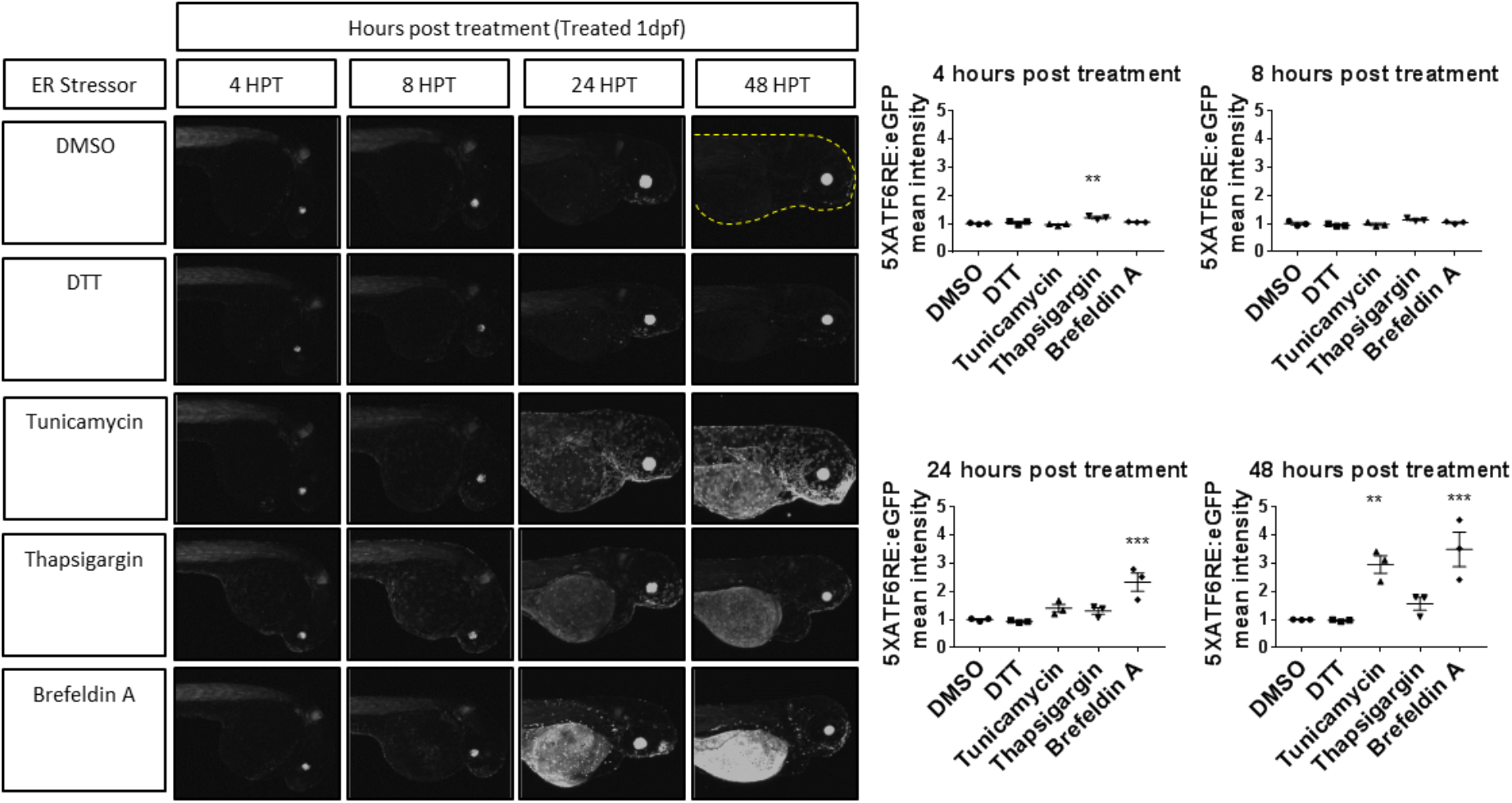
Chemical ER stressors activate eGFP reporter expression. (A) Zebrafish embryos were treated with ER stressors at 1dpf. Quantification (dotted line) revealed that *5XATF6RE*:eGFP expression was significantly higher at 4hpt with Thapsigargin (p=0.0012) treatment, 24hpt with Brefeldin A (p=0.0008) treatment, and 48hpt with Tunicamycin (p=0.0054) and Brefeldin A (p=0.0010) treatment. ** p≤0.01; *** p≤0.001; unpaired one-way ANOVA with Dunnett’s post-hoc test with respect to DMSO control group.

**Table S1:** Parent and final plasmids

| Plasmid Name | Parent Plasmid | Final Plasmid |
| --- | --- | --- |
| ATF6 constitutive active (caATF6) (1-373) | Adgene plasmid 27173, pCGN-ATF6 (1-373) | Tol2-UAS: caATF6 (1-373)-2a-mcherry |
| ATF6 dominant negative (dnATF6) (171-373) | Adgene plasmid 27173, pCGN-ATF6 (1-373) | Tol2-UAS: dnATF6 (171-373)-2a- mcherry |
| ATF4 full length | Adgene plasmid 82190, pDonr223_ATF4_WT | Tol2-UAS: ATF4-2a-mcherry |
| Spliced XBP1 (XBP1s) | Adgene plasmid 63680, pCMV5-Flag-XBP1s | Tol2-UAS: XBP1s-2a-mcherry |

